# Effects of SARS-CoV-2 Mutations on Protein Structures and Intraviral Protein-Protein Interactions

**DOI:** 10.1101/2020.08.15.241349

**Authors:** Siqi Wu, Chang Tian, Panpan Liu, Dongjie Guo, Wei Zheng, Xiaoqiang Huang, Yang Zhang, Lijun Liu

## Abstract

Since 2019, severe acute respiratory syndrome coronavirus 2 (SARS-CoV-2) causing coronavirus disease 2019 (COVID-19) has infected ten millions of people across the globe, and massive mutations in virus genome have occurred during the rapid spread of this novel coronavirus. Variance in protein sequence might lead to change in protein structure and interaction, then further affect the viral physiological characteristics, which could bring tremendous influence on the pandemic. In this study, we investigated 18 non-synonymous mutations in SARS-CoV-2 genome which incidence rates were all ≥1% as of July 15^th^, 2020, then modeled the mutated protein structures and compared them with the reference ones. The results showed that four types of mutations could cause dramatic changes in protein structures (RMSD ≥5.0 Å), which were Q57H and G251V in open reading frames 3a (ORF3a), S194L and R203K/G204R in nucleocapsid (N). Next, we found that these mutations could affect the binding affinity of intraviral protein interactions. In addition, the hot spots within these docking complexes were altered, among which the mutation Q57H was involved in both Orf3a-Orf8 and Orf3a-S protein interactions. Besides, these mutations were widely distributed all over the world, and their occurrences fluctuated as time went on. Notably, the incidences of R203K/G204R in N and Q57H in Orf3a were both over 50% in some countries. Overall, our findings suggest that SARS-CoV-2 mutations can change viral protein structure, binding affinity and hot spots of the interface, thereby may have impacts on SARS-CoV-2 transmission, diagnosis and treatment of COVID-19.

## Introduction

The ongoing outbreak of coronavirus disease 2019 (COVID-19), caused by severe acute respiratory syndrome coronavirus 2 (SARS-CoV-2), has been characterized as a pandemic by the World Health Organization (WHO). As of July 15^th^, 2020, there were a total of 13,150,645 confirmed cases and 574,464 deaths all over the world. SARS-CoV-2 is a positive-sense single-stranded RNA virus and belongs to Betacoronavirus [1], which genome is comprised of ~30,000 nucleotides, containing twelve open reading frames (ORFs) encoding four structural and twenty-two non-structural proteins. The structural proteins include spike protein (S), envelope protein (E), membrane protein (M), and nucleocapsid protein (N), while the non-structural proteins (nsp) contain nsp1~16 encoded by ORF1ab, and six accessory proteins which are Orf3a, 6, 7a, 7b, 8, and 9b [2]. It is reported that the evolution rate of a typical RNA virus is about 10^−4^ substitutes per year per site [3], and mutations could occur during each replication cycle. Published data also suggested that single nucleotide variants (SNVs) in SARS-CoV-2 genome were quite abundant. Hence, it would be important to find out all the mutations with relatively high incidences and investigate their potential impact on virus characteristics.

Non-synonymous mutations cause amino acid substitutions, and could alter virus protein structures, which may affect viral reproduction, leading to false-negative diagnoses and drug resistance. It is reported that the depletion of nine amino acids in SARS-CoV-2 Orf6 resulted in a dramatic alteration of protein structure and caused a shift in its transmembrane localization, which would lead to drug resistance of interferon (IFN) [4]. Additionally, along with the occurrence of S139A and F140A mutations, the structure of SARS-CoV 3CLpro altered remarkably, followed by the decrease of its enzyme activity [5]. Furthermore, alteration of protein structure by mutations also can affect protein-protein binding, and intraviral protein-protein interactions are indispensable in assembly and release of coronavirus. In SARS-CoV, the interactions between structural proteins are essential for its maturation [6], while the binding between non-structural proteins guarantees the completion of virus replication [7]. Y195A mutation in SARS-CoV M was found to disrupt its interaction with S, resulting in a declined ability of virus assembly [8]. However, it is still not clear whether some SARS-CoV-2 mutations lead to the changes of protein structures and protein-protein interactions, which could affect protein function, virus infection, and even clinical antiviral strategy. Hence, it is very important and urgent to find out SARS-CoV-2 mutations, especially the ones with high occurrence rate, that may result in the changes of virus characteristics.

In this report, we selected non-synonymous mutations of SARS-CoV-2 with ≥1% incidence, predicted and compared the structures of their corresponding proteins, and found that four types of mutations had significant impacts on protein structures, which resulted in remarkable changes in binding affinities and hot spots between virus proteins. Besides, statistical analyses on mutation rates exhibited dynamic changes in the process of time, and also demonstrated these four SARS-CoV-2 variants were widely distributed and had relatively high incidences in certain countries. Thus, our findings on SARS-CoV-2 mutations would be very helpful for better understandings about this virus and dealing with the complicated situations in COVID-19 prevention, diagnosis and treatment.

## Results

### SARS-CoV-2 mutations led to altered protein structures

As the COVID-19 pandemic is spreading around the world, thousands of SARS-CoV-2 mutations have been evolved. To investigate these SARS-CoV-2 nucleotide polymorphisms, first we took one of the earliest reported SARS-CoV-2 genome sequences (GenBank accession number: MN908947.3) as a control. Next, from CNCB 2019nCoVR database (see Methods), we selected 18 mutations with frequency ≥1% among all non-synonymous mutations, as of July 15^th^, 2020, which were located in ten different SARS-CoV-2 protein-coding regions (Table S1). Specifically, there was one amino acid substitution in nsp2, nsp3, nsp5, nsp6 or S. Meanwhile, two single substitutions were observed in nsp12, nsp13, Orf3a or Orf8, among which mutation P323L in nsp12 had the highest incidence (41.62%), and four single ones were found in N. In particular, there were also some combined mutations, such as P504L/Y541C in nsp13, and R203K/G204R in N (Table S1). Briefly, there were 15 types of mutants composed of 18 mutations with ≥1% incidence, which included one nsp2, nsp3, nsp5, nsp6, nsp13 or S mutant, two nsp12, Orf3 or Orf8 mutants, and three N mutants.

To examine whether these mutations had effects on protein structure, we predicted three-dimensional (3D) structures of each mutant and its reference protein using I-TASSER (Table S1) [9]. The structural similarity between mutant and control protein were compared using TM-align [10]. After structural alignment, we found that four mutants exhibited significant difference in protein structural morphology from their control ones (RMSD ≥5.0 Å), which were Q57H Orf3a, G251V Orf3a, S194L N, R203K/G204R N, respectively (Fig. 1 and Table S1).

**Figure 1.**
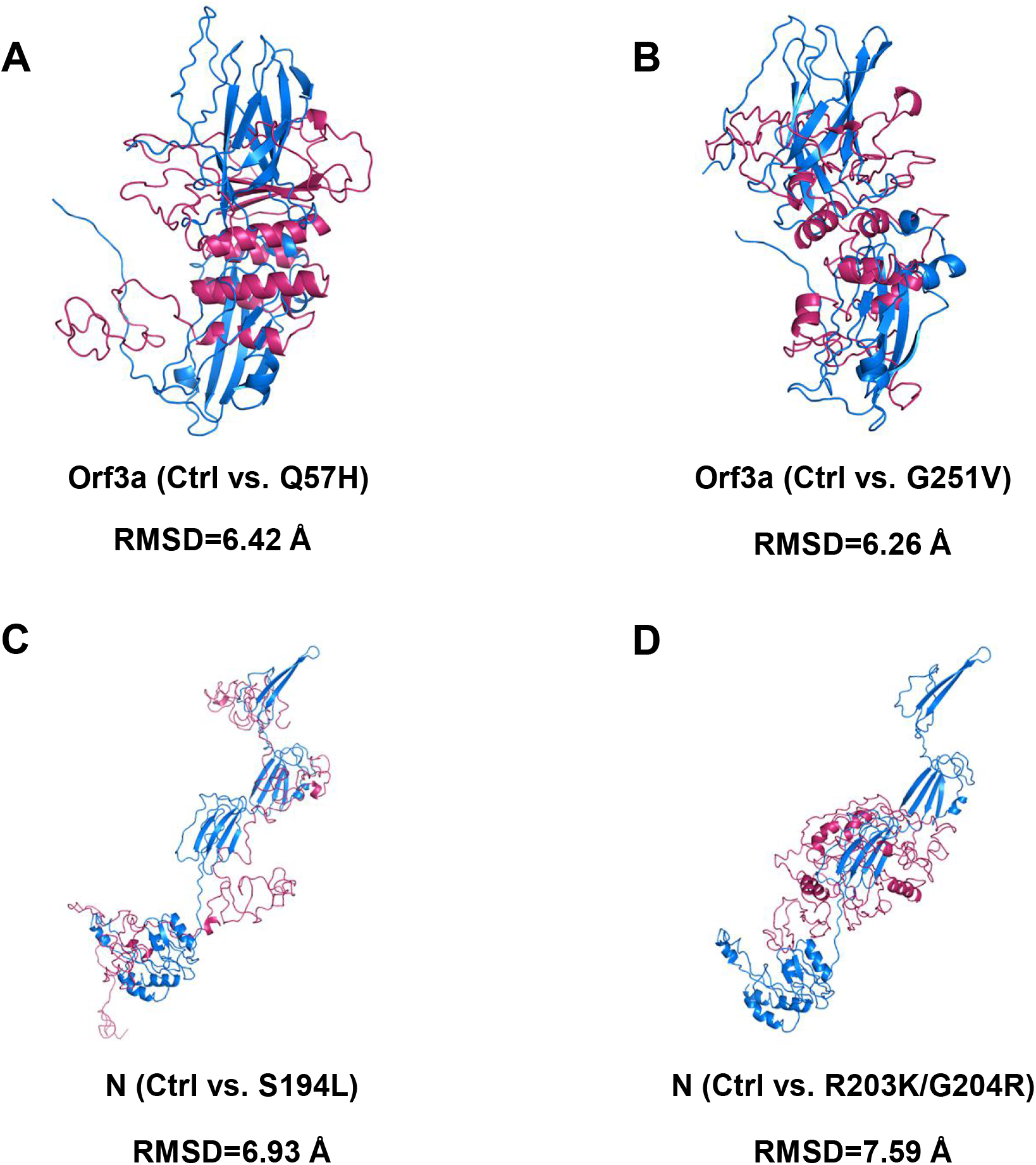
Comparisons of protein structures between SARS-CoV-2 control and mutated proteins. The aligned structures of control ones and Q57H Orf3a (A), G251V Orf3a (B), S194L N (C) and R203K/G204R N (D) are shown in blue (Ctrl) and warm pink (mutant) with the value of RMSD below.

### SARS-CoV-2 mutations resulted in the changes in protein-protein interactions

The SARS-CoV-2 proteins have been shown to display characteristic SARS-CoV features [11, 12]. So here, we referenced SARS-CoV protein combinations and explored the effect of these mutations which caused the alterations of protein structure on intraviral SARS-CoV-2 protein interaction known as a rate-limited procedure for virus reproduction [13–16]. Among 16 mutant docking pairs, five mutant complexes had significantly higher binding affinity than control ones (Fig. 2). Differently, another four mutant pairs showed weakened binding affinity in comparison with the control, and there was no obvious change in the remaining seven groups which absolute values of ΔΔG (= ΔG_control_ − ΔG_mutant_) were less than 1 kcal/mol (Fig. 2). Strikingly, binding affinity of Q57H Orf3a-S complex showed the greatest increased (ΔΔG = 4.2 kcal/mol), while the most dramatic decrease was observed in binding affinity of G251V Orf3a-M complex (ΔΔG = −2.3 kcal/mol) (Fig. 2A). Moreover, in the process of Orf3a interacting with M or S, Q57H mutant pairs showed enhanced binding affinity, while G251V mutant pairs had attenuated ones as just mentioned (Fig. 2A), suggesting that diverse amino acid substitutions in the same protein could lead to different effect on binding affinity.

**Figure 2.**
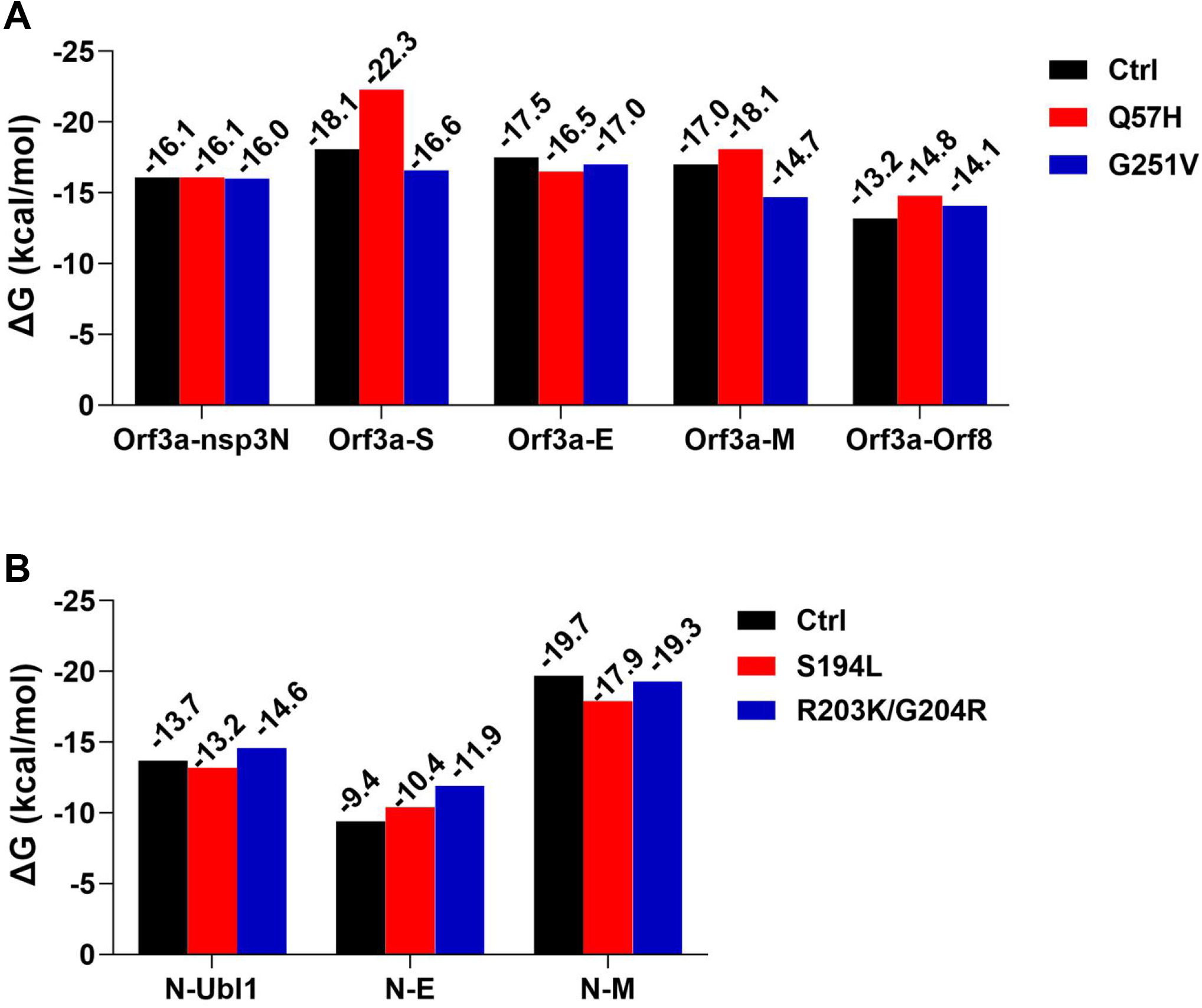
Protein binding affinity in SARS-CoV-2 mutated complexes. Bars show Gibbs free energy (ΔG) for representing binding affinity in Orf3a (A) and N (B) relevant complexes. Nsp3N: 1-595aa of nsp3, Ubl1: 1-112aa of nsp3.

### SARS-CoV-2 mutations caused varied hot spots within protein complexes

Hot spots are functional sites within protein-interacting interfaces, which are conservative and often taken as attractive drug targets via preventing protein-protein interactions. To further study the influence of SARS-CoV-2 mutations on protein binding hot spots, we predicted the hot spots in nine mutated complexes with altered bind affinity using the KFC2 server (Figs. 3, 4 and S1). Surprisingly, the results showed that the binding hot spots on mutated protein-protein interfaces were notably different from control ones, and few identical hot spots were shared by both the mutant and control complexes. In particular, amino acid substitution Q57H was a hot spot in Orf3a-Orf8 and Orf3a-S complexes, but Q57 residue in control Orf3a was not involved in the protein binding interfaces (Figs. 3B and 4A). These results indicated that SARS-CoV-2 mutations might destroy drug targeting sites and lead to therapy failure by shifting protein-binding interface.

**Figure 3.**
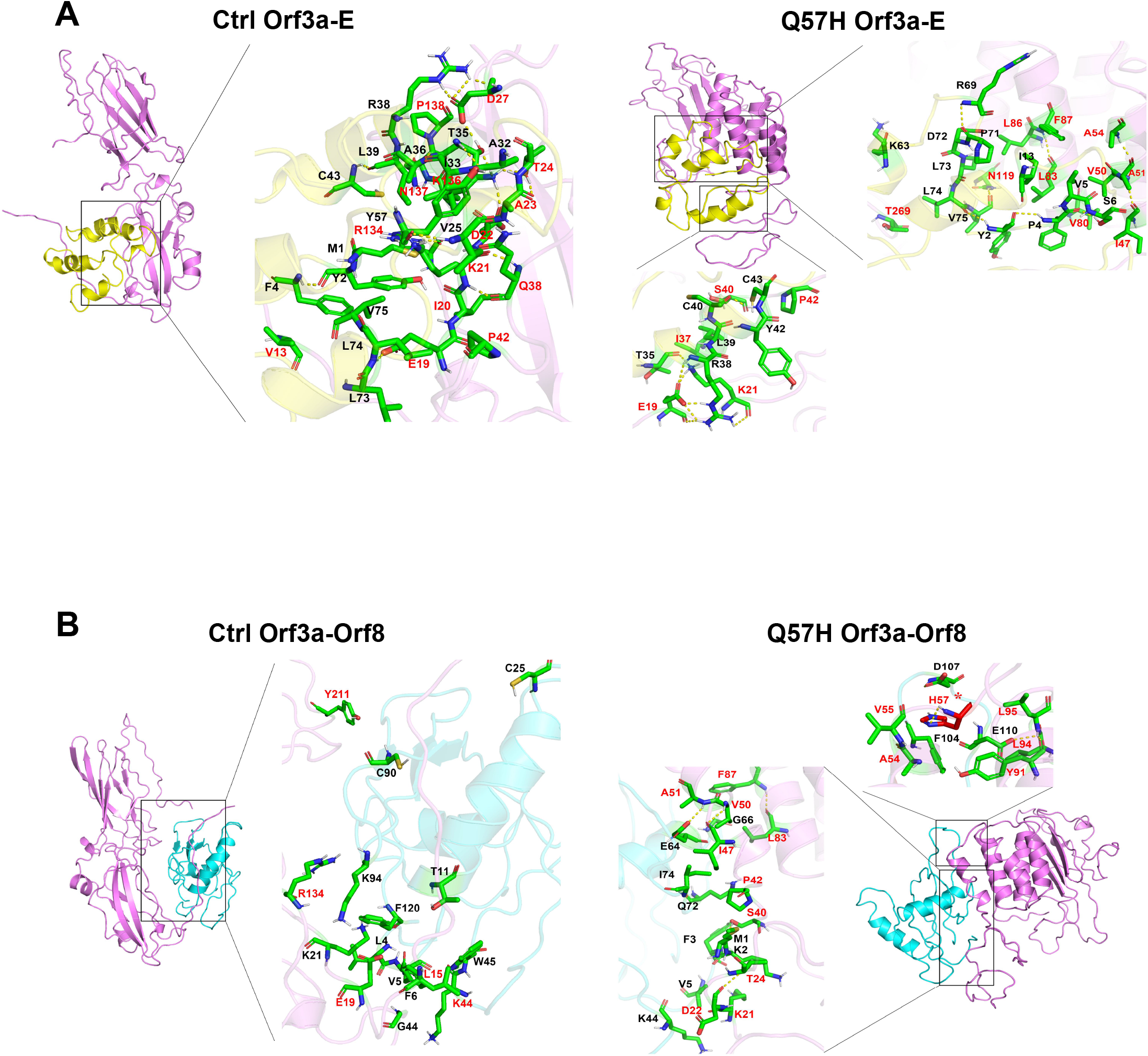
Hot spots within interactions between SARS-CoV-2 Orf3a and E or Orf8. The hot spots within interactions between Orf3a (Ctrl or Q57H) and E (A) or Orf8 (B) are shown as sticks. Control and mutated Orf3a are shown in violet, and E or Orf8 is shown in yellow or cyan. The residues of Orf3a and docking proteins are colored in red and black, respectively. Asterisk (*) represents mutated residue.

**Figure 4.**
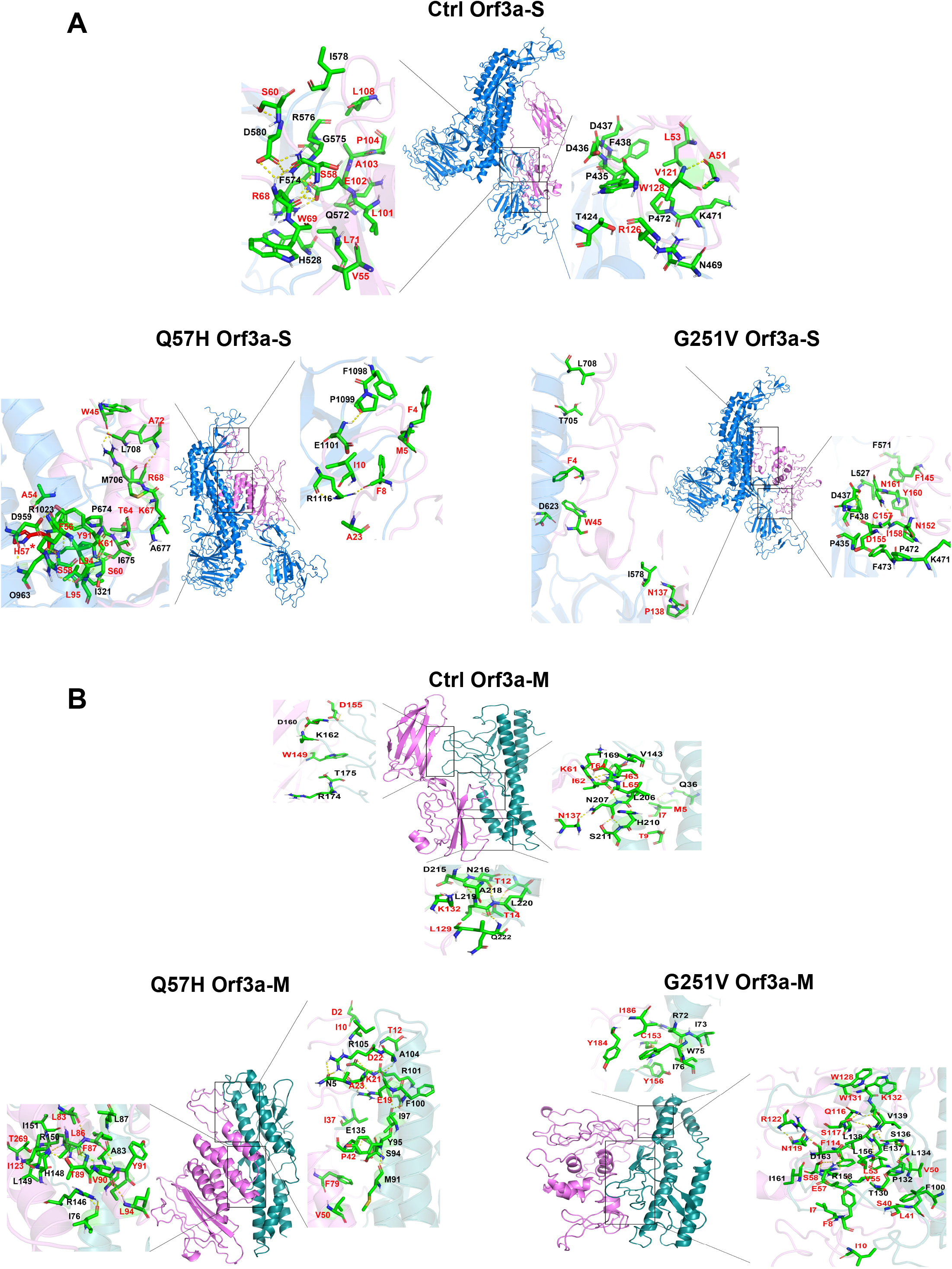
Hot spots within interactions between SARS-CoV-2 Orf3a and S or M. The hot spots within interactions between Orf3a (Ctrl, Q57H or G251V) and S (A) or M (B) are shown as sticks. Control and mutated Orf3a are shown in violet, and S or M is shown in marine or deep teal. The residues of Orf3a and docking proteins are colored in red and black, respectively. Asterisk (*) represents mutated residue.

### SARS-CoV-2 mutations were globally distributed with dynamic incidences over time

The changes in SARS-CoV-2 proteins caused by mutations can affect virus transmission, pathogenesis and immunogenicity. And with the spread of this pandemic, the incidence and lethality of SARS-CoV-2 infection varied from country to country. Hence, to link SARS-CoV-2 mutations and COVID-19 prevalence for controlling the pandemic, we analyzed the occurrence of these four types of mutations (Q57H and G251V in Orf3a, S194L and R203K/ G204R in N) based on CNCB 2019nCoVR database. To avoid deviation caused by insufficient sample size, we screened out the countries or regions where the number of SARS-CoV-2 genome sequences submitted was no more than 100, as of July 15^th^, 2020. The statistical results showed that high mutated incidence areas were distributed all over the world except Antarctica with the dynamic occurrences over time (Fig. 5). The frequency of all these mutations were relatively high in some countries (Fig. 5). Among them, we noticed that the incidence of R203K/G204R in N was as high as 85.44% in Bengal, meanwhile, it was also over 50% in Russia or Greece, with a trend of escalation (Fig. 5D). Analogously, the frequency of Q57H in Orf3a in Saudi Arabia, Finland and Denmark was shown the high incidences, which were 73.38%, 69.4% and 58.97% respectively (Fig. 5A).

**Figure 5.**
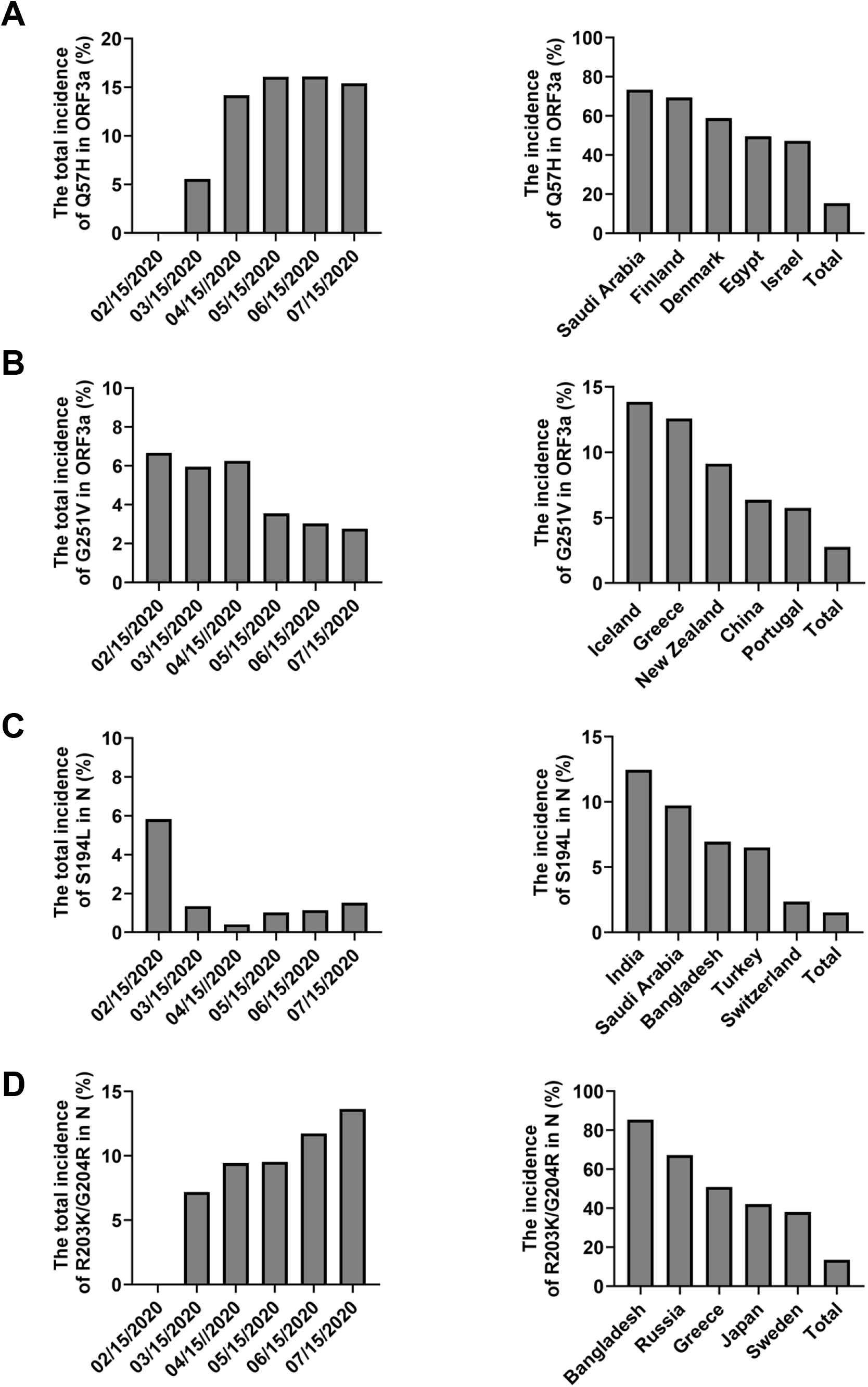
Temporal and spatial analyses of SARS-CoV-2 mutation incidences. Bars show the incidences of Q57H (A) and G251V (B) in Orf3a, S194L (C) and R203K/G204R (D) in N at different time periods or in top five countries and globe (Total), as of July 15^th^, 2020.

## Discussion

With rapid dissemination of SARS-CoV-2, thousands of single nucleotide polymorphisms (SNPs) have been evolved and widely distributed. According to CNCB 2019nCoVR database as of July 15^th^, 2020, we screened 18 non-synonymous mutations with ≥1% incidence, which were categorized into 15 types of mutation combinations. After structural alignment, four mutated proteins (Q57H Orf3a, G251V Orf3a, S194L N and R203K/G204R N) were found displaying different structures from control N and Orf3a that were essential for coronavirus assembly [17] and cytotoxicity [18]. Consistent with our data, some predicted results showed that R203K/G204R in N also resulted in destabilized protein structure [19]. But there were some mutations, such as T85I in nsp2 and P323L in nsp12, found no significant impact on integral protein structure according to our results. However, their high frequency prompt us to further investigate whether they could affect virus characteristics via altering RNA second structure [20], protein stability [21] or partial structure [22]. For instance, experimental research reported that D614G in S could lead to virus infectivity increased by eliminating side-chain hydrogen bond which was only a tiny change in overall protein structure [23], and our data also showed small structural difference between D614G mutant and control (RMSD = 2.33 Å). Therefore, more research about SARS-CoV-2 mutations are worth doing to determine whether structure alterations of Orf3a or N would affect protein function and even virus infectivity.

Besides protein structure alteration, the mutations in virus genome could also affect protein-protein interactions. With molecular docking and ΔG value calculation, we found that protein binding affinity changed between all four above mutants and their docking intraviral proteins. It has been confirmed that these protein interactions in SARS-CoV play indispensable mediating roles in virus characteristics [6, 18, 24], thus their changes may affect its infection. Moreover, the combination of SARS-CoV Orf3a and S can prevent the release of premature viral RNA [25], and Orf3a-M complex is localized in Golgi [26], where virus particles are assembled by budding [27]. Our results showed stronger binding affinities in Q57H Orf3a-M and Q57H Orf3a-S complexes, but weaker affinities in G251V Orf3a-M and G251V Orf3a-S complexes. Considering the incidences of Q57H in Orf3a (15.40%) and G251V in Orf3a (2.77%) showed a great disparity, we speculated that Orf3a mutants might be relevant to virus assembly and transmission. Furthermore, some researches indicated that the interaction between N and E had a function in SARS-CoV release [15, 28, 29], and the N-M complex was necessary for the assembly of coronavirus [16]. Therefore, the enhanced interaction between each SARS-CoV-2 N mutant (S194L or R203K/G204R) and E might promote virus release, while decreased binding affinity of S194L N-M might attenuate virus assembly. Besides, SARS-CoV N can bind to heterogeneous nuclear ribonucleoprotein A1 (hnRNP A1) of host cells and their interaction play a regulatory role in the synthesis of SARS-CoV RNA [30], so it would be interesting to check the combination of these N mutants and the proteins in host cells. What is more, the hot spots play critical roles in protein-protein interactions, which were generally used as drugs target, so their changes could have great impacts on clinical treatment [31]. Our results demonstrated that all these selected SARS-CoV-2 mutations (Q57H and G251V in Orf3a, S194L and R203K/G204R in N) had great influences on hot spots within protein combinations, which indicated that it is essential to take into account the interference of SARS-CoV-2 mutations during the development of vaccines or drugs [32].

In the last several months, residents in more than 180 countries/regions have been affected by SARS-CoV-2, and the emergence of mutations might already make important contributions to virus adaption to new environments and selective pressures, which in turn can impact transmissibility, pathogenesis and immunogenicity of SARS-CoV-2 [33]. It has been reported that variant SARS-CoV-2 genomes occurred in different areas [34–38]. Our results also showed that some mutations were widely distributed, and their occurrence rates had different dynamic fluctuations. Take R203K/G204R in N as an example, this mutation combination with increased incidence was found mainly in Europe till March 2020 [39, 40], while our data showed its high frequency was also in Asian countries as of July 2020, suggesting that these two mutations might be conducive to SARS-CoV-2 adaptability and spread. Therefore, it is particularly important to pay attention to the difficulties in COVID-19 diagnosis and treatment that probably caused by these mutations, especially in countries with high occurrence rates. Although there is still no reliable evidence for any necessary link between mutations and epidemic outbreak in specific countries/regions, it is urgent to identify the relevance of geographically aggregated mutations to SARS-CoV-2 transmission and pathogenicity for effective containment of COVID-19 outbreaks.

## Methods

### Online resources

All information about the mutated SARS-CoV-2 genomes were obtained from China National Center for Bioinformation 2019 Novel Coronavirus Resource (CNCB 2019nCoVR) (https://bigd.big.ac.cn/ncov). Data were visualized by GraphPad Prism 8.

### Protein structure prediction

Protein structure of control SARS-CoV-2 S was from C-I-TASSER (https://zhanglab.ccmb.med.umich.edu/C-I-TASSER/2019-nCoV/) [41], while other SARS-CoV-2 protein structure models were predicted by I-TASSER [9]. For each protein, five models were generated and the model with the highest C-score was selected as the best one and used for the following analysis.

### Protein structure alignment

The mutated protein structures were aligned to corresponding control ones by using the TM-align web-server [10]. Random structural similarity was determined by TM-score between 0.0~0.3 and Root-mean-square deviation (RMSD) ≥5.0 Å [10, 42].

### Molecular docking and hot spots prediction

Protein-protein docking was performed with the HADDOCK web-server (http://haddock.chem.uu.nl/) [43]. The structure was chosen according to the HADDOCK score, and complex binding affinity was calculated by PRODIGY [44, 45]. Hot spots within protein-protein interfaces were predicted using Knowledge-based FADE and Contacts Server (KFC, https://mitchell-web.ornl.gov/KFC_Server/index.php) [46]. Data visualization was accomplished by PyMOL.

## Supplemental Information

Supplemental information includes one figure and one table.

## Acknowledgments

We thank Xiaoting Li (Columbia University, USA) for helpful advice on modeling analysis. And we also would like to thank Dr. Qi Zhao (Northeastern University, China) for technical assistance on molecular docking. This study was supported by Northeastern University, PRC (N2020006), and Talent Project of Revitalizing Liaoning (XLYC1907052). The funders had no role in study design, data collection and analysis, decision to publish, or preparation of the manuscript.

## Author Contributions

S.W., C.T. and L.L. conceived the study. S.W., C.T. and P.L. collected and summarized data of SARS-CoV-2 mutations information. Y.Z., W.Z., C.T., S.W. and P.L. contributed to protein structure prediction. X.H., C.T., S.W. and P.L. contributed to molecular docking. C.T., S.W. and P.L. predicted hot spots within the protein-protein interface. D.G. contributed to protein structures alignment and data visualizations. S.W., C.T. and L.L. wrote the paper. L.L. directed the study.

---

**Figure S1. Hot spots within interactions between SARS-CoV-2 N and E or M**

The hot spots in N (Ctrl, S194L or R203K/G204R)-E (A), and N (Ctrl or S194L)-M (B) complexes are shown as sticks. Control and mutated N are shown in grey, and E or M is shown in yellow or deep teal. The residues of N and docking proteins are colored in red and black, respectively.

**Table S1. Information of SARS-CoV-2 mutations and protein models**

