## Supplementary material for "Effects of SARS-CoV-2 Mutations on Protein Structures and Intraviral Protein-Protein Interactions": Figure S1 and Table S1

A

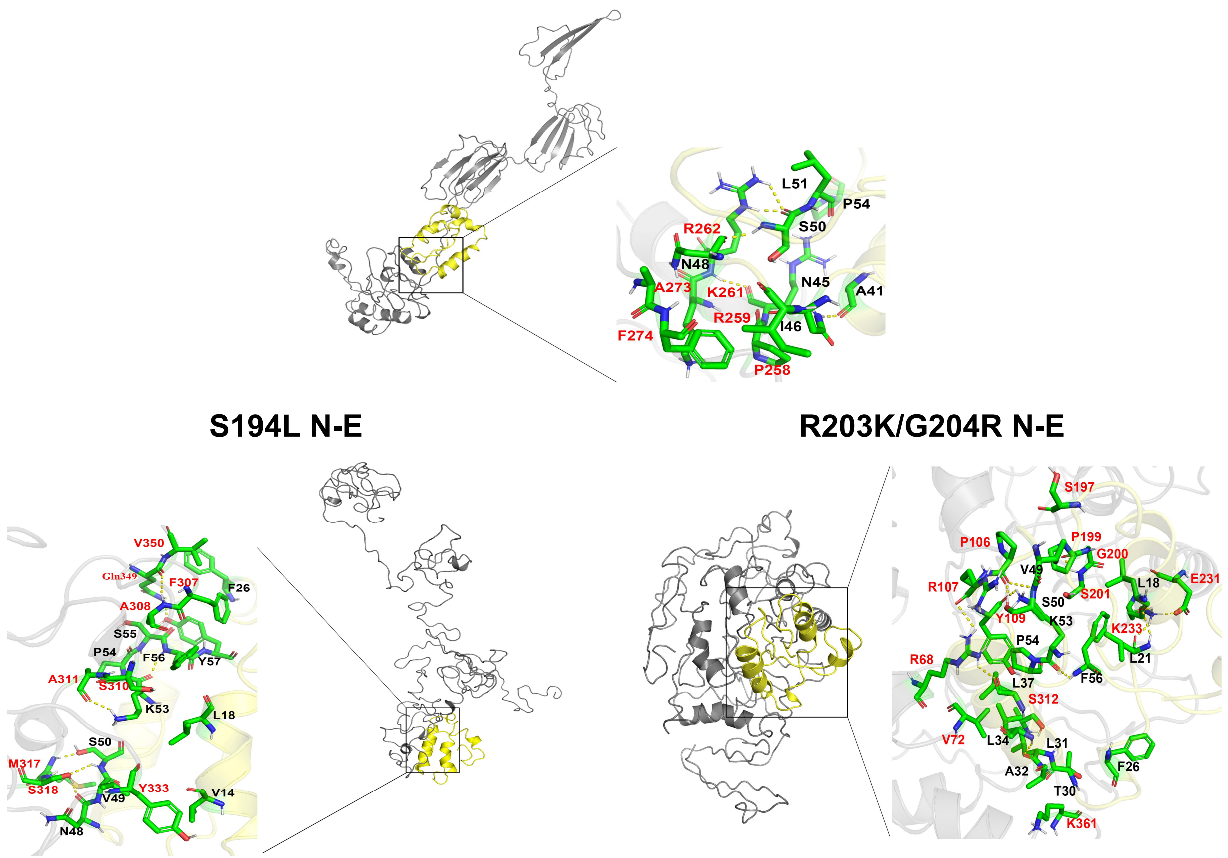
Ctrl N-E
L51
P54
R262
S50
N48
N45
A41
A273
K261
R259
I46
F274
P258
S194L N-E
V350
Gln349
F307
F26
A308
S55
P54
F56
Y57
A311
S310
K53
L18
S50
M317
S318
V49
Y333
V14
N48
R203K/G204R N-E
S197
P106
P199
G200
V49
R107
S50
S201
L18
E231
Y109
K53
K233
P54
R68
L21
F56
L37
S312
V72
L34
L31
F26
A32
T30
K361

B

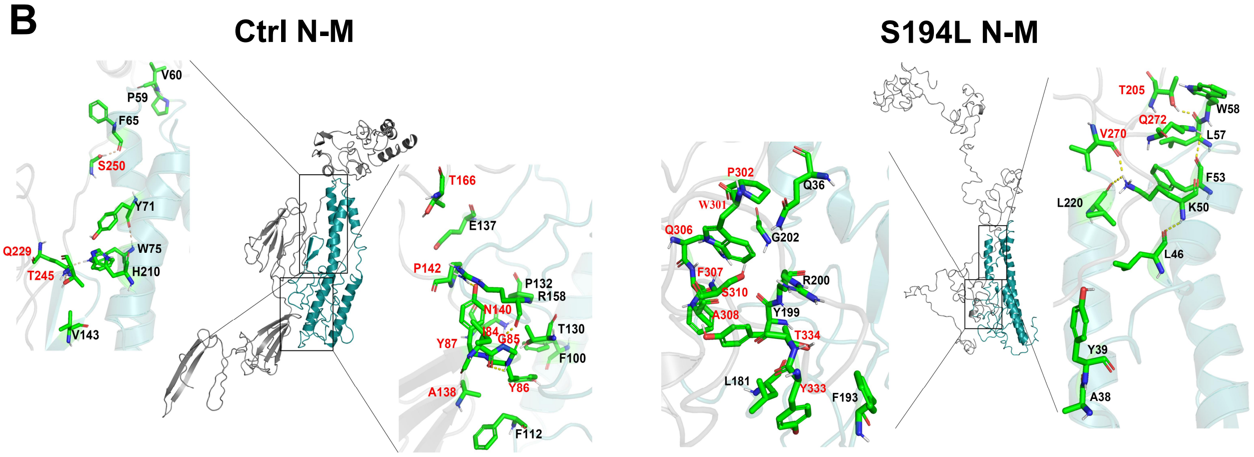
Ctrl N-M
V60
P59
F65
S250
Y71
Q229
W75
T245
H210
V143
T166
E137
P142
P132
R158
N140
I84
G85
T130
Y87
F100
Y86
A138
F112
S194L N-M
P302
Q36
W301
Q306
G202
F307
R200
S310
Y199
A308
T334
L181
Y333
F193
T205
W58
Q272
V270
L57
F53
L220
K50
L46
Y39
A38

**Table S1. Information of SARS-CoV-2 mutations and protein models**

| Gene name/ Region | Base substitution(s) | Amino acid substitution(s) | Incidence | C-score | Estimated TM-score | TM-score (TM-align) | RMSD (TM-align) |
|---|---|---|---|---|---|---|---|
| nsp2 | N/A | - | - | -1.48 | 0.53 | - | - |
| | C1059T | T85I | 0.12 | -1.57 | 0.52 | 0.9330 | 1.17 |
| nsp3 | N/A | - | - | -1.75 | 0.5 | - | - |
| | C6312A | T1198K | 0.0103 | -1.72 | 0.51 | 0.9354 | 2.37 |
| nsp5 | N/A | - | - | 1.84 | 0.97 | - | - |
| | G10097A | G15S | 0.0145 | 1.86 | 0.98 | 0.9984 | 0.27 |
| nsp6 | N/A | - | - | -2.14 | 0.46 | - | - |
| | G11083T | L37F | 0.0504 | -2.07 | 0.47 | 0.8721 | 2.55 |
| nsp12 | N/A | - | - | -1.30 | 0.55 | - | - |
| | C13730T | A97V | 0.0111 | -0.87 | 0.6 | 0.8703 | 1.08 |
| | C14408T | P323L | 0.4162 | -0.85 | 0.61 | 0.8709 | 1.39 |
| nsp13 | N/A | - | - | 2.00 | 0.99 | - | - |
| | C17747T/ A17858G | P504L/Y541C | 0.0251 | 2.00 | 0.99 | 0.9992 | 0.24 |
| S | A23403G | D614G | 0.4019 | -1.40 | 0.54 | 0.8879 | 2.33 |
| ORF3a | N/A | - | - | -3.14 | 0.36 | - | - |
| | G25563T | Q57H | 0.1540 | -3.47 | 0.33 | 0.2244 | 6.42 |
| | G26144T | G251V | 0.0277 | -4.38 | 0.26 | 0.2350 | 6.26 |
| ORF8 | N/A | - | - | -3.88 | 0.3 | - | - |
| | C27964T | S24L | 0.0148 | -3.59 | 0.32 | 0.7080 | 3.12 |
| | T28144C | L84S | 0.0482 | -3.99 | 0.29 | 0.7689 | 2.67 |
| N | N/A | - | - | -1.58 | 0.52 | - | - |
| | C28311T | P13L | 0.0110 | -1.62 | 0.52 | 0.9585 | 2.08 |
| | C28854T | S194L | 0.0154 | -0.34 | 0.67 | 0.2452 | 6.93 |
| | GGG28881-28883AAC | R203K/G204R | 0.1363 | -1.59 | 0.52 | 0.1841 | 7.59 |

Note: The data about SARS-CoV-2 mutations was collected from CNCB 2019nCoVR on July 15$^{th}$, 2020.
